# Large ungulates will be present in most of Japan by 2050 owing to natural expansion and human population shrinkage

**DOI:** 10.1101/2024.01.23.576970

**Authors:** Takahiro Morosawa, Hayato Iijima, Tomonori Kawamoto, Takahisa Kanno, Ryota Araki, Teruki Oka

## Abstract

The aims of this study were to elucidate factors contributing to the expansion of the distributions of sika deer and wild boar in Japan and to predict the expansion of their distributions by 2025, 2050, and 2100. A site occupancy model was constructed using information on species distribution collected by the Ministry of the Environment in 1978, 2003, 2014, and 2018, days of snow cover, forested and road areas, elevation, human population, and distance from occupied grid cells as covariates to calculate the probability of distribution change. Factors contributing to distribution expansion were elucidated and distribution expansion was predicted. Distance from occupied grid cells had the strongest influence on distribution expansion, followed by the inherent ability of each species to expand its distribution. For sika deer, human population had a strong negative effect and elevation and number of days of snow cover were important. For wild boar, forest area and elevation had high importance. Predictions of future distribution showed that both species will be distributed over 90% of Japan by 2050 and over 100% by 2100.

## Introduction

Large ungulates have both positive and negative effects on human life. They offer services of hunting (Schulp et al. 2014), venison (Goguen et al. 2017), and medicine (Wu et al. 2013) and disservices of the irreversible alteration of ecosystems by over-browsing (Royo and Carson 2011; Tanentzap et al. 2012), economic damage in forestry and agriculture (Cote et al. 2004), collisions with vehicles (Gunson et al. 2011), and the transmission of zoonoses (Hale et al. 2022) and of disease vectors such as ticks (Ostfeld et al. 2018; Iijima et al. 2022). Planning for future human well-being depends on the prediction of population expansion of large ungulates and related factors.

The distribution of large ungulates has continued to expand in recent years, especially in developed countries, and it is urgent to understand why (Carden et al. 2011; Sales et al. 2017; Cretois et al. 2021; Iijima et al. 2023). Factors related to their expansion include climate change (Ohashi et al. 2016), land use change (Acevedo et al. 2011; Carpio et al. 2021), and hunting (Keuling et al. 2008; Cromsigt et al. 2013; Linnell et al. 2020; Iijima et al. 2023). Because global climate change is expected to continue, it may accelerate the expansion of their distribution (Ohashi et al. 2016; Markov et al. 2019). The relationship between climate and distribution has been related to snow (Markov 1997; Kaji et al. 2000; Honda 2009; Månsson et al. 2012), and it is expected that as snowfall decreases with climate change, species distributions will expand into more northern and heavier-snowfall regions. In addition, human populations are shrinking in developed countries, so associated changes in land use, such as abandonment of land (Acevedo et al. 2010) and reduced hunting pressure (Acevedo et al. 2011; Tsunoda and Enari 2020), may accelerate expansion. Therefore, it is important to predict distribution expansion and plan responses in regions where the distributions of large ungulates are expected to expand.

To predict the expansion of the distribution of large ungulates, researchers have used species distribution models (SDMs; e.g., Markov et al. 2019; Rutten et al. 2019) and models that focus on the process of distribution expansion (e.g., Zurell et al. 2009; Acevedo et al. 2014; Morelle and Lejeune 2015). SDMs are powerful tools for predicting the expansion of distribution and explaining the relative importance of environmental factors related to expansion (Elith and Leathwick 2009). Environmental factors in newly occupied areas can differ from those in original habitats, yet animals adapt to the new resources (Mainali et al. 2015). Thus, spatial information is also an important aspect of distribution prediction and including this information should lead to higher predictive power of SDMs.

As both the likelihood of distribution expansion and the similarity of environmental factors depend on distance, it is important to consider spatial information in relation to both. Incorporating spatial information such as distance from the current distribution into models while taking environmental factors into account will improve the realism of predictions. In addition, while adding temporal information allows for better predictions, the computational load is even greater. Therefore, it is necessary to consider ways to reduce the computational load when estimating temporal dynamics in large-scale spatial-scale predictions.

In Japan, large-scale land use changes have occurred since the period of rapid economic growth with expanded afforestation, population growth, and urbanization began in the 1960s (Ministry of the Environment 2006). Under these changes, the populations of large ungulates such as sika deer and wild boar have recovered under national protection policies (Kaji et al. 2000; Yamazaki et al. 2016). Both species were distributed throughout Honshu in the prehistoric Jōmon period, but their distributions had been shrinking since the Meiji period (1868–1912) (Tsujino et al. 2010). They expanded again during the period of expanded afforestation after World War II (Iijima et al. 2023). Since then, the distributions of both species have continued to increase, sika deer by 2.7 times in area and wild boar by 1.9 times from 1978 to 2018 (Ministry of the Environment 2021). In 2018, both species were reported to have expanded into the northern Tōhoku and Hokuriku regions, where they had not lived since the end of World War II owing to heavy snowfall (Ministry of the Environment 2021). With such dynamic changes in landscape and wildlife distribution and its wide latitudinal gradient, Japan is a suitable region in which to evaluate factors associated with changes in wildlife distribution.

The main aim of this study was to clarify factors important for population expansion of large ungulates. We also predicted the distribution of both sika deer and wild boar up to 2100. Climate change and human population shrinkage (Enari 2021) are also occurring in many other developed countries. Thus, prediction of the distribution of both species by considering the important factors is necessary in order to understand these species’ future expansion.

## Materials and methods

### Data set of sika deer and wild boar distribution

We used data collected by the Ministry of the Environment’s Natural Environment Survey in 1978 and 2003 from wildlife keepers and local government to map the distribution of sika deer and wild boar (http://gis.biodic.go.jp/webgis/?_ga=2.193090862.2011052859.1653966617-587115043.1623632179). Data for 2014 were collected as areas where the distributions had expanded since 2003 as reported by local government. Data for 2014 and 2018 were provided by the Office of Wildlife Management, Ministry of the Environment. All data were collected and used on a 5-km × 5-km scale.

### Explanatory variables

#### Physical environmental variables

Average altitude, land use, and road area in each cell were drawn from national land numerical information (https://nlftp.mlit.go.jp/ksj/, 1-km × 1-km grid cell). The elevations recorded in 2009 were used for all years and averaged 1-km × 1-km grid cell data including in each 5-km × 5-km grid cell.. Land use data in 1976, 2006, and 2015, which approximate the years of animal data, were also drawn from national land numerical information, from which areas of forest, agricultural land, paddy fields, water, and building land were extracted. Road area data in 1978, 2002, and 2010 were similarly drawn. Forest area and road area of the 1-km x 1-km grid cells were summed up with the 5-km x 5-km grid cells to create a data set.

#### Climate variables

We used data on temperature and total rainfall from the meteorological stations of the Japan Meteorological Agency (JMA; https://www.data.jma.go.jp/obd/stats/etrn/) and calculated the averages. Although the JMA has about 1300 meteorological stations nationwide, there are some areas where data are not available. Therefore, we interpolated missing values by spatial complementation of the values of other meteorological stations by Delaunay triangulation on a 5-km × 5-km scale. We used data on the deepest snow depth and the number of snow days provided by Ohashi and Kominami .

#### Human population

We used national census data (1 km × 1 km; https://www.e-stat.go.jp/) from 1980, 2005, and 2015. The data of 1-km x 1-km grid cells contained in a 5-km x 5-km grid cells was summed up.

#### Distance from cells in which ungulates are present

Distances from cells in which sika deer and wild boar are present were calculated for 1978, 2003, 2014, and 2018 in GIS (ArcGIS v. 10.5). We calculated the center of each cell from the 5-km grid data and calculated the distances between centers. In the case of cells in which a species is present, the distance is 0 km; for a cell in which the species is absent, the distance from the nearest occupied cell was calculated.

#### Data for future prediction

For future prediction, we used the above physical environment data, meteorological data, human population, distance from occupied cells, and the 2009 elevation data. For land use, road area, and human population, we used the results predicted by the National Institute for Environmental Studies’ “S8” project (https://www.nies.go.jp/s8_project/). In the “S8” project’s predictions were made in multiple scenarios, but values with median sea level were used in this study. For meteorological data, we used predictions made by the Ministry of the Environment with the MRI-NHRCM20 regional climate model at a 20-km scale, using the same values for each 5-km × 5-km cell within each larger cell, and the median SST2 as the sea surface temperature. Ten-year average weather data were used in consideration of the uncertainty of the predictions. In predicting future distributions, we used the RCP2.6 climate change scenario, which has the lowest temperature rise, and RCP8.5, which has the highest temperature rise, to compare the effects of climate change.

### Model construction

It is common to use statistical models, such as SDM, to estimate spatial distributions of species (Phillips et al. 2006). Such models assume that the distribution of animals is defined by environmental factors at each location. It is common that spatially closer locations may have similar distributions, which is explained by spatial autocorrelation. Distributions cannot be properly predicted without consideration of spatial autocorrelation (Legendre and Legendre 1998), and tools for doing so include generalized linear mixed models (GLMMs), conditional autoregressive GLMMs, conditional autoregressive models (CARs), and simultaneous autoregressive models (SARs) (Keitt et al. 2002; Dormann 2007; Dormann et al. 2007). The latter two methods impose a high computational load and are therefore difficult to apply to a wide-scale time-series analysis such as ours. In GLMMs, data that are considered to be not independent can be divided into blocks, and the effects of spatial autocorrelation can be considered simply by treating the blocks as random effects, but the subjective division of blocks can create arbitrariness. The method of weighting explanatory variables according to distance has also been used to consider spatial autocorrelation (Dormann et al. 2007). However, we were interested in whether the ability of sika deer and wild boar to expand their distributions or the environmental conditions had a stronger effect, so if we weighted the variables by distance, we would not be able to properly evaluate the effect on the ability to expand distribution.

In constructing a model for the expansion of the species’ distribution, we first considered a model in which the distribution probability is explained only by environmental factors. However, strong spatial autocorrelation in spatially close environments may lead to incorrect predictions if spatial autocorrelation is not taken into account (Legendre and Legendre 1998). For this reason, it was necessary to consider spatial autocorrelation as well. However, it was difficult to do so here because of the high computational load due to the large number of cells (16L023) and the prediction of time-series distribution expansion. Therefore, we decided instead to add the distance from an occupied cell as an explanatory variable. This method is appropriate when the factors causing spatial autocorrelation are biologically clear (Miller et al. 2007). As the purpose of this study was to evaluate how the ability of a species to expand its distribution affects the actual distribution, we consider this an appropriate method.

The use of statistical models to estimate potential habitat is common in predicting distributions (e.g., Phillips et al. 2006). However, problems can include not considering changes over time and overestimating the environmental effect of a yet-unoccupied or newly occupied area (Pearman et al. 2008). Considering temporal change as here allows us to evaluate the selectivity of the environment by taking into account the process of dispersion, making predictions more realistic.

Mammals in particular have a strong ability to disperse, and the process of dispersal, or its spatial parameters, is considered to be more important than environment in predicting distribution. Large mammals move seasonally (e.g., Takii et al. 2012; Morelle and Lejeune 2015), and home ranges change by a factor of 2 seasonally (Morelle and Lejeune 2015). Thus, unless certain environmental factors strongly limit distribution, mobility or dispersal ability contributes more strongly than environment to the expansion of the distribution. Thus, it is necessary to consider both movement and dispersion ability and spatial autocorrelation when predicting distribution (Månsson et al. 2012). However, when we try to consider spatial autocorrelation over such a large area as here, the calculation load can be prohibitively large. Therefore, we used distance from the current distribution as easy-to-manage spatial information to improve prediction accuracy.

To construct the distribution expansion model, we used a site occupancy model (MacKenzie et al. 2002; Kéry and Royle 2016) as a hierarchical model based explicitly on an observation model that explains the probability of distribution in each cell in terms of environmental factors and distance from an occupied cell, and a process model that determines the presence or absence of distribution in each cell (Royle and Dorazio 2008). The site occupancy model is based on both the probability that presence/absence at a site will continue unchanged and the probability that a site will be newly occupied. We constructed a distribution prediction model using the species distribution data and environmental factor data prepared above. First, we tested the correlations between the factors by Spearman’s rank correlation, and for those with ρ ≥ 0.6, we used either one. We found a strong correlation between the number of snow days and temperature; snow cover contributes significantly to the distribution of both species (Tokita et al. 1980; Maruyama 1981). We selected the number of snow days.

The dynamics of a target species is expressed by the process model as:

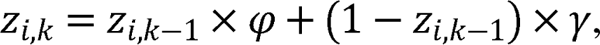

where *z_i,k_* is the probability of the presence of a target species in the *i*th cell in the *k*th year; φ is the probability of survival; and γ is the probability of establishment. The equation predicts the presence/absence of the species in each cell from the sum of these probabilities. As the species will continue to live in places where they already live and will not become absent unless there is a great deal of trouble, we expressed this situation only by the probability of new establishment:

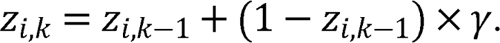

As the initial distribution, we assumed a uniform Bernoulli distribution with expected values of 0 to 1, and used random numbers to correspond to the distribution data.

We expanded the process model by incorporating covariates into γ. The value of γ in each cell will be affected by the environment, land use, meteorological factors, human population, and distance from an occupied cell, so we expressed it as:

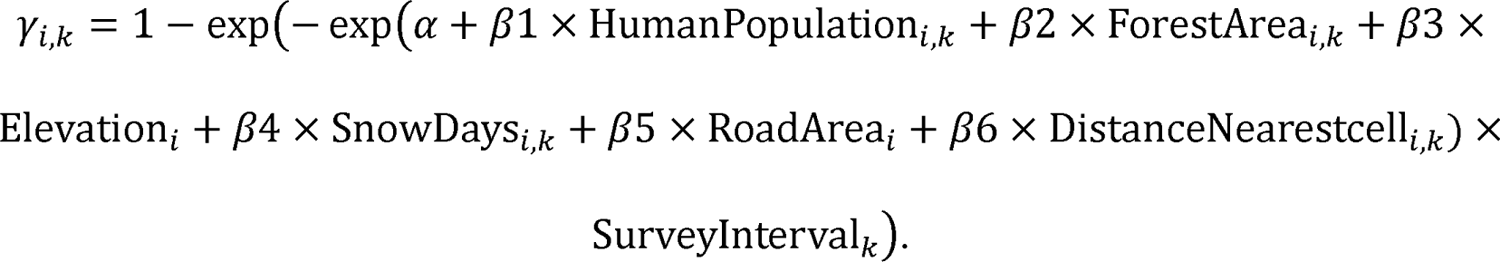

Using the complementary log–log function as above allows us consider the effect of the difference between survey intervals on the presence/absence of wildlife. The function takes into account the variation in intervals between survey years, which are used as an offset term. In addition, as the interval of the year for which the distribution data is obtained is not equal, we used the interval of the year predicted by including the predicted interval of the year as the change in the distribution per unit time. As the coefficients of each factor, we set a normal distribution (0, 1000) as the prior distribution. In addition, since the large number of cells to which factor data were only partially applied stymied the estimation, we used the method of estimating the factor itself from the data and substituting it. The factor was calculated as a normal distribution (expected value of factor, variance of factor), and its variance as a hyperparameter with a uniform distribution (0, 100). All explanatory variables were standardized before analysis, so the relative importance of the coefficients could be evaluated from their magnitude by comparing the estimates. A Markov Chain Monte Carlo model was run with a burn-in of 50L000 steps, a calculation 200L000 steps, and thinning of 15. We judged that the calculated R̂ converged below 1.1 (Gelman et al. 2013). The calculation was performed by the JAGS v. 4.3.0 tool (Plummer 2017) in R v. 4.02 (R Development Core Team 2020).

## Results

### Factors that contribute to distribution expansion

The distance from occupied cells had the greatest effect on the expansion of the distributions of both species, followed by the number of snow days and forest area (Table 1). For sika deer, forest area, snow days, and road area had positive effects, whereas human population, elevation, and distance from distribution had negative effects. For wild boar, forest area and road area had positive effects, whereas human population, elevation, snow days, and distance from distribution had negative effects. Only the effect of snow days differed in sign between species. In sika deer, magnitudes of coefficients decreased in the order of distance from occupied cells > snow days = forest area > road area > elevation > human population. In wild boar, they decreased in the order of distance from the occupied cells > forest area > elevation > number of snow days > human population > road area. The importance of distance from occupied cells is also indicated by the fact that the coefficient of snow days for both species is greater for the model that did not use distance than for the model that used distance (Supplement Table 1).

**Table 1.** Estimated coefficients of predictive variables in each species model with distance.

| Variable | Deer | Wild boar |
| --- | --- | --- |
| Intercept | −15.1 (−15.52 to −14.77) | −62.61 (−66.97 to −36.27) |
| Population | −0.84 (−1.09 to −0.62) | −0.14 (−0.22 to −0.08) |
| Forest area | 0.36 (0.30 to 0.41) | 0.58 (0.51 to 0.76) |
| Elevation | −0.25 (−0.30 to −0.19) | −0.21 (−0.28 to −0.14) |
| Snow days | 0.36 (0.30 to 0.41) | −0.20 (−0.43 to −0.09) |
| Road area | 0.23 (0.12 to 0.33) | 0.07 (0.02 to 0.11) |
| Distance from occupied cell | −23.43 (−24.04 to −22.93) | −94.73 (−101.28 to −55.04) |

### Distribution prediction

In the distribution prediction models for sika deer (Fig. 1) and wild boar (Fig. 2) under the RCP2.6 scenario, the correct response rate within the time range for which distribution data were available (i.e., 1978 to 2020) was 100%, confirming the model’s high predictive power. Therefore, we calculated results for 2025, 2050, and 2100. The results show that sika deer will be present in 99.6% of the cells by 2050 and 100% by 2100 (Fig. 3), with wild boar in 93.8% of the cells by 2050 and 100% by 2100 (Fig. 4).

**Fig. 1.**
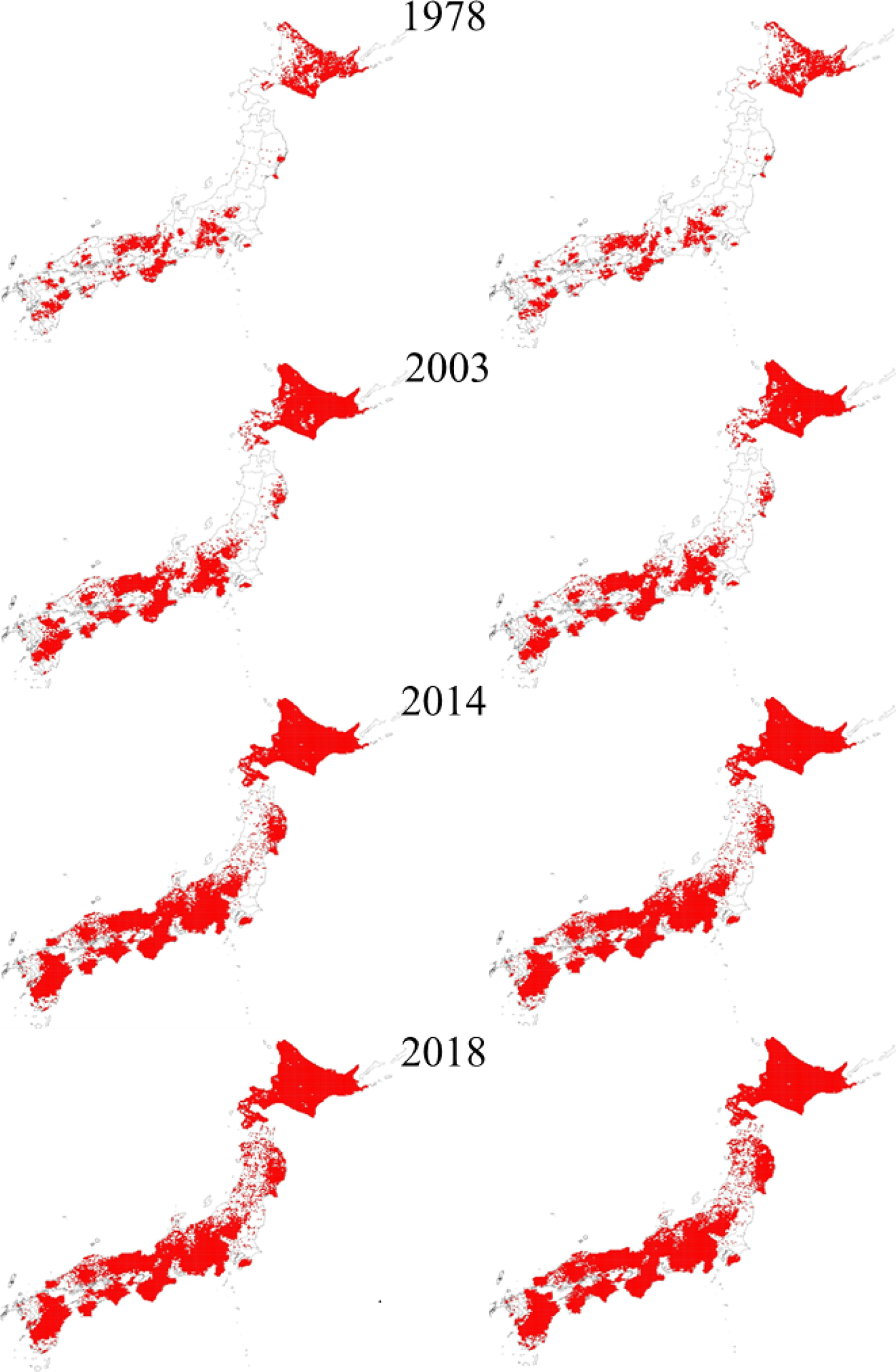
Model training data (left column) and predictions (right column) for sika deer under the RCP2.6 scenario.

**Fig. 2.**
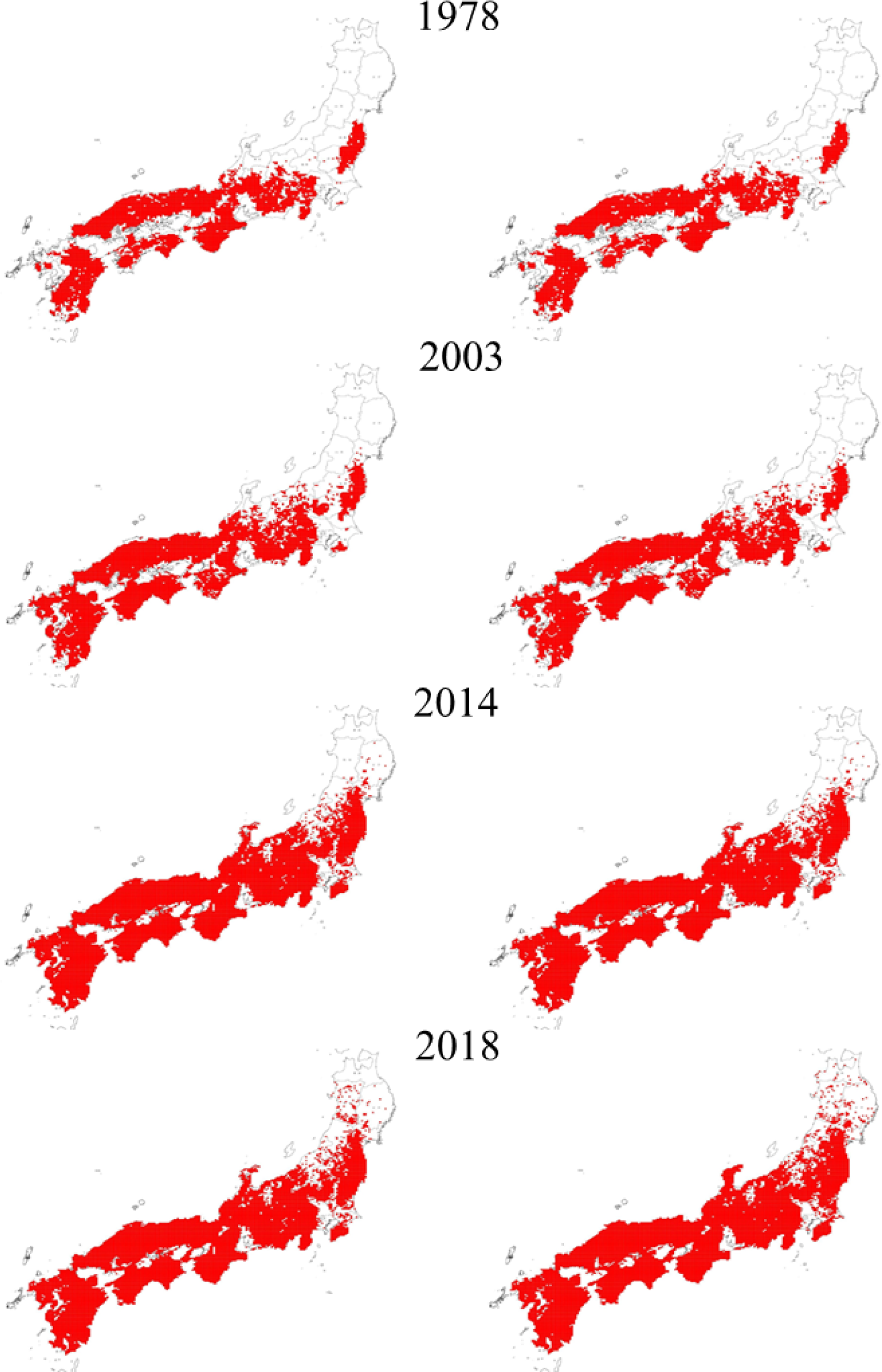
Model training data (left column) and predictions (right column) for wild boar under the RCP2.6 scenario.

**Fig. 3.**
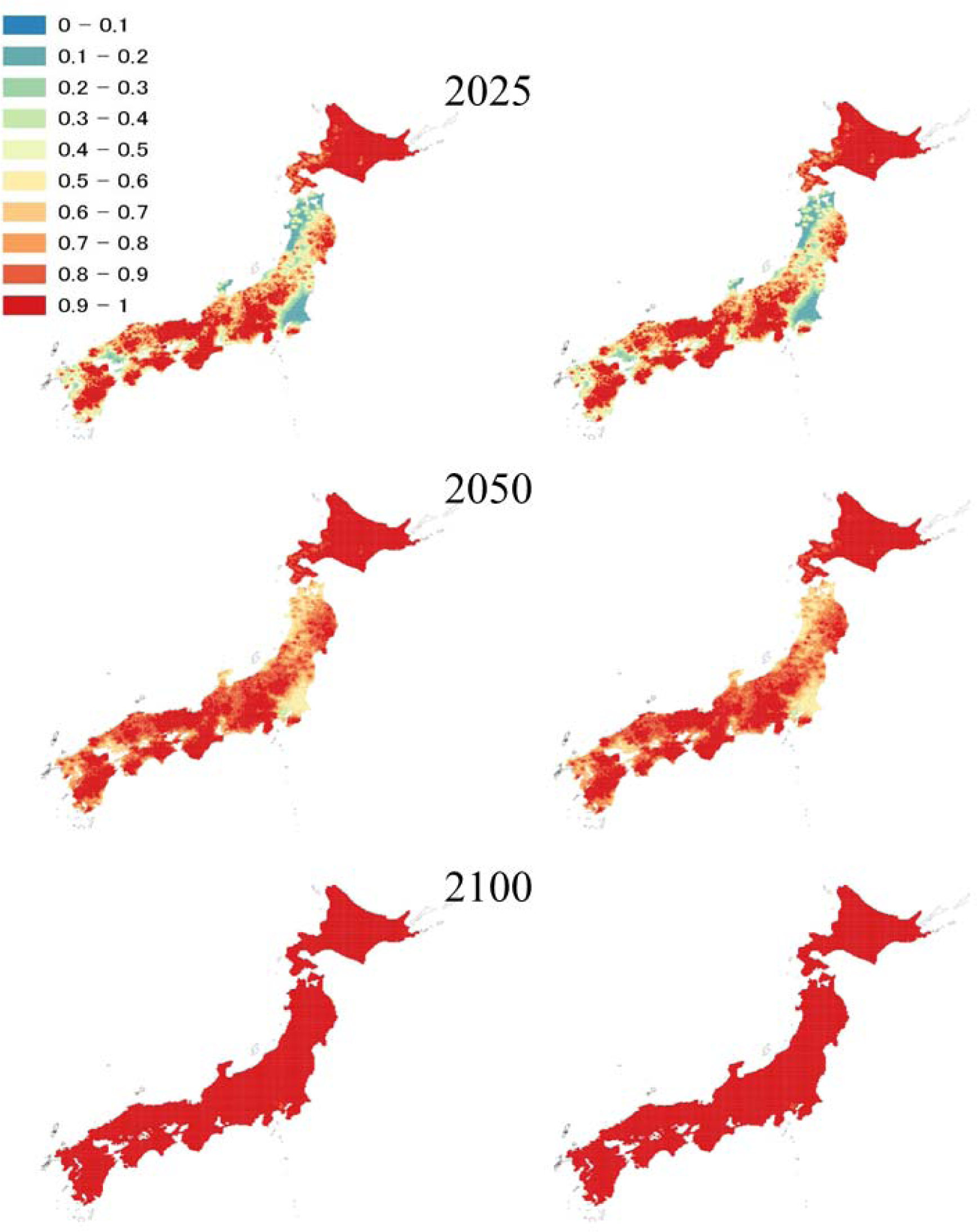
Predicted probabilities of deer distributions in 2025, 2050, and 2100 under the RCP2.6 (left column) and RCP8.5 (right column) climate change scenarios.

**Fig. 4.**
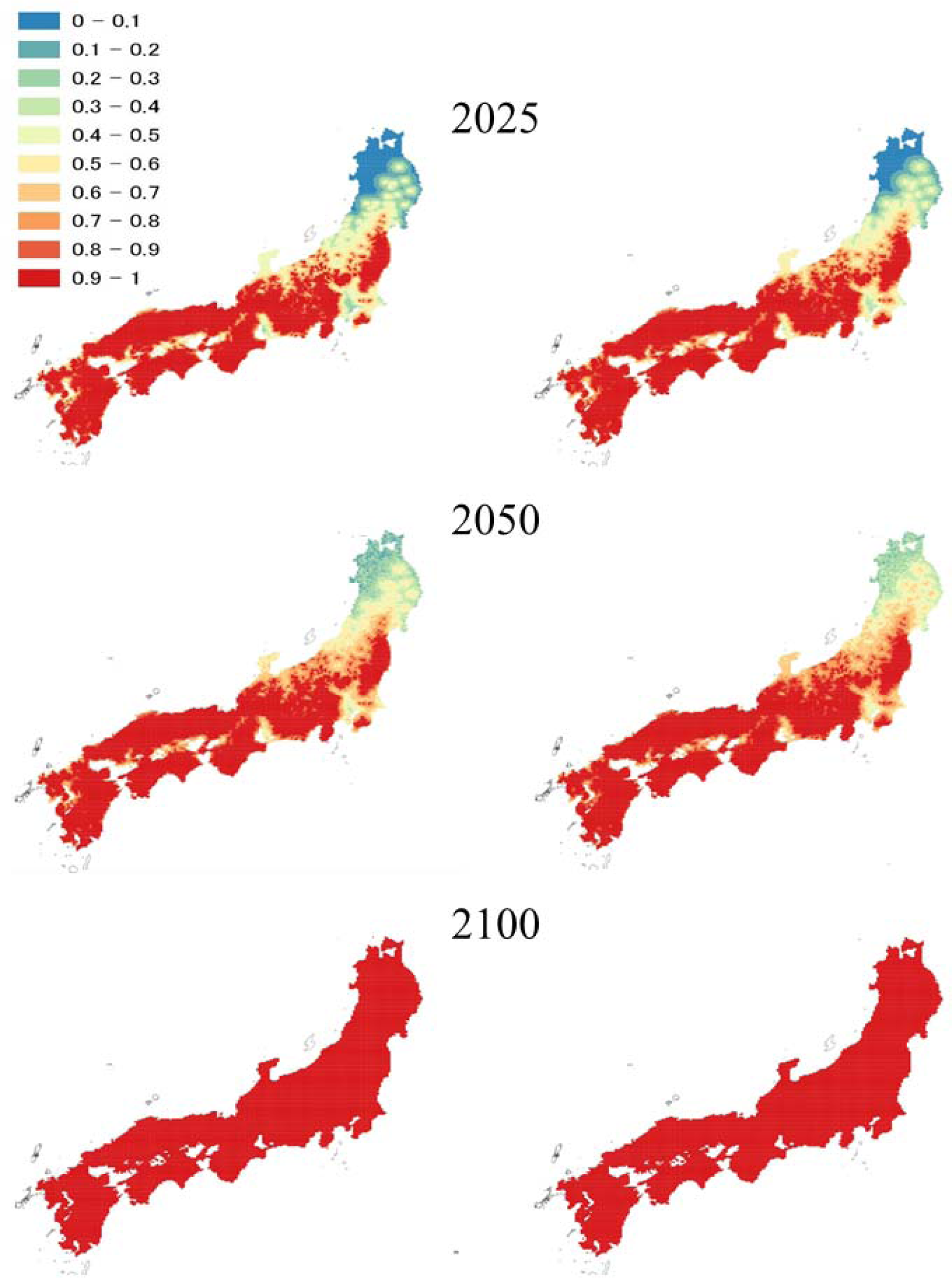
Predicted probabilities of wild boar distributions in 2025, 2050, and 2100 under the RCP2.6 (left column) and RCP8.5 (right column) climate change scenarios.

Under RCP2.6 and RCP8.5, the distributions of both species are expected to expand throughout Honshu by 2050, with an estimated probability of almost 100% in 2100 (Figs. 3, 4). The results of both species are very similar between scenarios. However, the slightly higher distribution probability in 2050 under RCP8.5 indicates that climate change may accelerate the distribution expansion of wild boar in particular.

## Discussion

### Factors that contribute to the expansion of distribution

We identified the relative importance of factors that contributed to the expansion of sika deer and wild boar populations between 1978 and 2018 and their differences between the species (Table 1). By using distance from occupied cells as spatial information, we could examine the effects of climatic factors, human population, and land use on the expansion of distributions. By comparing the species, we obtained new findings.

As previous studies repeatedly showed, snowfall was one of the important factors defining the distribution of large ungulates (Kaji et al. 2000; Okumura et al. 2009; Ohashi et al. 2016; Saito et al. 2016; Markov et al. 2019). Although snow days affected the population expansion of both species, its effect was positive on deer but negative on wild boar. Our results suggest that the distribution of sika deer is likely to expand in the future in areas where snow depth is still high. Recent reports indicate that coniferous forests, which tolerate snow cover, function as wintering grounds for sika deer (Igota et al. 2004; Takii et al. 2012), which can be seen in large numbers there even in areas with heavy snowfall, where sika deer have not historically been seen (Kaji et al. 2000), supporting our results. Our contrasting results for wild boar reflect their shorter legs: deep snow will continue to suppress their distribution, but climate change may facilitate their expansion. Wild boar have been observed in polar regions recently, so they can survive in areas with heavy snow and low temperatures (Markov et al. 2019).

Distributions of both species did not differ greatly between the RCP2.6 and RCP8.5 scenarios. Thus, the number of snow days contributed somewhat to their distributions, but much less than the distance from occupied cells and human population. Thus, temperature increase due to climate change may slightly accelerate the distribution expansion.

Human population had a relatively high negative effect on distribution expansion. The expansion of the distribution of deer species is expected to be restricted in urban areas (Anderson et al. 2011) and areas with high hunting pressure (Saїd et al. 2012). On the other hand, behavioral flexibility allows wild boar to find suitable habitat in human-dominated landscapes (Rutten et al. 2019). Thus, the influence of areas with greater human conflicts may depend on species. For wild boar in particular, Stillfried et al. (2017) reported strong tolerance of human presence in urban areas, which our results support.

Forest area had a positive effect, because it functions as a migration route (Anderson et al. 2011; Okumura et al. 2009) and feeding area (Miyashita et al. 2008; Morelle and Lejeune 2015) for both species. Road area too had a positive effect, as dispersal or movement routes and environments provide feeding areas. In fact, both sika deer and wild boar have been reported to use roads for movement and feeding (Gerhardt et al. 2013; Miyashita et al. 2008). Especially on the Sea of Japan side of the country and in the Tōhoku region, where distribution has been expanding in recent years (Ministry of the Environment 2021), roads may serve as dispersal pathways to expand distribution.

### Results of distribution prediction

We predicted distributions in 2025, 2050, and 2100 from data collected in 1978, 2003, 2014, and 2018. The distributions of both species are still expanding, mainly on the Sea of Japan side of the country and in the Tōhoku region, but our results expand the distributions almost nationwide by 2050. Similarly, Ohashi et al. (2016) included human population trends as an explanatory variable and made their prediction in consideration of the influence of human population changes on the distribution of sika deer and wild boar.

Our prediction results take climate change and other factors into account. However, classical swine fever (CSF) has begun to spread among wild boar in recent years in Japan (Isoda et al. 2020), and an unexpected outbreak of infectious disease in sika deer cannot be ruled out. In addition, the Covid-19 epidemic has altered human activities. So changes that were not taken into account are possible, and the results may differ from the predictions.

Our predictions show that both sika deer and wild boar will be distributed over 90% of Japan by 2050 and over 100% by 2100. In 2016, Ohashi et al. (2016) projected that sika deer will be distributed in about 70% of the country by 2103, but our results suggest a faster expansion. Our study presents the first results for wild boar in Japan. In areas where distribution will expand in the future, prediction of when they will arrive and become established will enable countermeasures to be planned. We consider out results to be very important from the viewpoint of damage prevention.

### Implications for management

Our results indicate that the distribution of sika deer and wild boar will expand due to possible future changes in land use, human population dynamics, climate change, and other factors. Human population decline is expected to continue in Japan, so the distribution expansion of both species is likely to accelerate.

Major countermeasures are population management by hunting and prevention of damage. Hunting pressure affects the distribution of large ungulates (Keuling et al. 2008; Cromsigt et al. 2013; Linnell et al. 2020). Thus, it may be possible to delay the expansion of distribution through hunting. Traditionally, hunting pressure has depended on local hunters, and strategic use has not been attempted in Japan. Our results will make it possible to examine, for example, where hunting pressure can be most effectively applied to delay expansion. In addition, since we clarified factors that are positively related to distribution expansion, it is possible to use our results to identify where hunting pressure should be intensively applied. Fences are set up after damage has occurred, but through the use of the predicted distributions, they can be set up in advance to prevent damage. In addition, our results can be used to prepare budgets in advance and to study damage control methods. Therefore, efficient and effective application of hunting pressure and damage control based on distribution prediction will become necessary in Japan.

## Acknowledgements

We thank Dr. Haruka Ohashi and Dr. Yuji Kominami for providing the snow data and the Office of Wildlife Management, Ministry of the Environment, for the sika deer and wild boar distribution data for 2014 and 2018.

## Declarations

### Ethics approval

This article does not contain any studies with human participants or animals performed by any of the authors.

### Consent to participate

Not applicable.

### Consent for publication

Not applicable.

### Availability of data and material

All datasets are available from the corresponding author on reasonable request.

### Code availability

The code used is available from the corresponding author on reasonable request.

