## Supplemental Table 1 for "Large ungulates will be present in most of Japan by 2050 owing to natural expansion and human population shrinkage"

### Supplement

Table 1 Estimated coefficients of predictive variables in each species model without distance.

| Variable | Deer | Wild boar |
|---|---|---|
| Intercept | −2.71 (−1.95 to −3.53) | −27.82 (−28.6 to −27.1) |
| Population | 5.69 (+6.40 to +5.03) | −1.64 (−2.12 to −1.17) |
| Forest area | 1.09 (+1.80 to +0.39) | 0.19 (−0.15 to +0.6) |
| Elevation | −1.33 (−0.58 to −2.1) | 8.76 (+8.45 to +9.11) |
| Snow days | 1.00 (+1.63 to +0.37) | −23.62 (−24.22 to −22.97) |
| Road area | −2.71(−1.95 to −3.53) | −0.45 (−0.91 to +0.056) |
